# *Listeria monocytogenes* biofilm-derived cells show differential *sig*B expression on a food model and enhanced survival in simulated gastric conditions

**DOI:** 10.64898/2026.04.27.721029

**Authors:** Raquel A. Nogueira, Juan J. Rodríguez-Herrera, Pedro Rodríguez-López, Marta L. Cabo

## Abstract

*Listeria monocytogenes* is a foodborne pathogen of utmost interest to food industry stakeholders because it persists in food processing environments. The ability to form biofilms – bacterial communities of autoaggregated cells embedded in a self-produced matrix – contributes to its persistence. While it is known that biofilm cells exhibit different gene expression than their planktonic counterparts, it remains to be elucidated whether those differences persist once cells detach from the biofilm and what their implications might be for food safety. Therefore, this study examines the differential *sig*B expression in biofilm-derived cells from three *L. monocytogenes* strains isolated from the environment within a food model subjected to varying osmotic stress over a 15-day storage period. Under our experimental conditions, biofilm-derived *L. monocytogenes* cells showed higher *sig*B expression compared to planktonic counterparts. The upregulation was strain-dependent and transient, suggesting that physiological memory may influence stress adaptation during early storage but dissipates over time. Then, the safety implications of *sig*B upregulation in biofilm-derived cells were assessed by evaluating cell survival under a simulated gastric environment (pH 1-3). The biofilm-derived cells showed a significant increase in survival under severe gastric conditions compared to the planktonic counterparts. Overall, our findings highlight the need to consider biofilm-derived cells in shelf-life studies and predictive models to more accurately reflect real contamination scenarios. Relying exclusively on planktonic cultures introduces a bias that may compromise risk analysis and decision-making.

## 1. Introduction

*Listeria monocytogenes* is a pathogen of utmost importance for being the aetiological agent of listeriosis, a severe foodborne disease with a mortality rate of about 20 % [1,2]. Despite the implementation of control measures across the food chain, from environmental sources of production systems to food shelf life, *L. monocytogenes* has shown a notable ability to survive and proliferate in food processing environments (FPE). The pathogen’s persistence in industrial settings poses a significant, ongoing challenge to global food safety. In particular, ready-to-eat (RTE) products are a primary concern for listeriosis risk because they are consumed without a heating process. The European Food Safety Authority (EFSA) report [3] indicates that approximately 90 % of invasive listeriosis cases result from consuming RTE products, according to quantitative microbial risk assessment (QMRA) models.

To survive in FPE and foods, *L. monocytogenes* undergoes complex physiological adaptations, driven by transcriptional networks that regulate stress response and persistence [4,5]. Among the molecular regulatory mechanisms driving environmental adaptation, the alternative sigma factor B (*sig*B) constitutes the largest regulon [6]. This factor drives the cellular response to major environmental stressors, such as cold [7], heat, oxidative stress, and osmotic or acidic conditions [8,9]. Additionally, *sig*B has been shown to be involved in biofilm formation [10].

The ability to form or integrate into surface-attached biofilms is a key survival strategy for *L. monocytogenes*, especially under environmental stress [11]. Biofilms are complex communities of attached microorganisms embedded in a self-produced extracellular polymeric matrix [12]. The biofilm state constitutes the predominant mode of life for microorganisms in nature. Biofilms confer resistance on bacteria, increasing their tolerance to environmental stressors and thereby enabling their recalcitrance, making them a source of contamination within FPE. Several studies have shown that *L. monocytogenes* biofilm cells exhibit significantly different gene expression profiles compared to the planktonic counterparts [13–15]. Specifically, mature biofilm cells have been observed to show increased *sig*B expression compared to their planktonic counterparts [10,16].

The *sig*B expression differences between biofilm and planktonic states reflect the community-wide shift to a high-resistance phenotype and may predispose cells to distinct stress responses upon transfer to foods. However, the influence of the cellular origin – either biofilm or planktonic – on the pathogen’s molecular and physiological responses remains largely unaccounted for in QMRA. This omission is significant because biofilm-derived cells, whether surface-attached or detached, may exhibit distinct survival characteristics when transferred into a food matrix compared with planktonic cells. Upon transfer to a food matrix, *L. monocytogenes* encounters a range of stresses, such as osmotic stress and low pH, where differential *sig*B activation in the previous state could enhance cell tolerance and adaptation. This issue is highly relevant to food contamination, as these physiological states may lead to distinct resistance phenotypes that could ultimately increase the risk of listeriosis.

Therefore, this study aims to explore differences in *sig*B expression and survival between *L. monocytogenes* cells from biofilms and planktonic cultures under food-storage and simulated-gastric conditions. It begins by analysing differential gene expression among three strains isolated from food and FPE, focusing on genes associated with persistence, motility, and virulence. Next, cells from biofilm and planktonic origins were transferred to a food model simulating the saline concentrations encountered in RTE products to assess differential *sig*B expression. Lastly, the fitness of biofilm- and planktonic-derived cells on the food model was assessed by evaluating their survival in a simulated gastric environment at different pH levels. Overall, this study provides new insights into the differences between biofilm- and planktonic-derived cells when exposed to osmotic and acid stress following an episode of cross-contamination.

## 2. Materials and Methods

### 2.1. Bacterial Strains and culture conditions

Three *L. monocytogenes* (**Table 1**) were used in this study. Stock cultures were preserved at - 80 °C in sterile brain-heart infusion broth (BHI; Scharlab, Barcelona, Spain) mixed 1:1 with 50 % (v/v) glycerol. Working cultures were stored at -20 °C. For strain reactivation, 100 µL of the corresponding working culture was transferred to 5 mL of Tryptone Soy Broth (TSB, Scharlab, Barcelona, Spain), followed by overnight incubation without shaking at 37 °C.

**Table 1.**
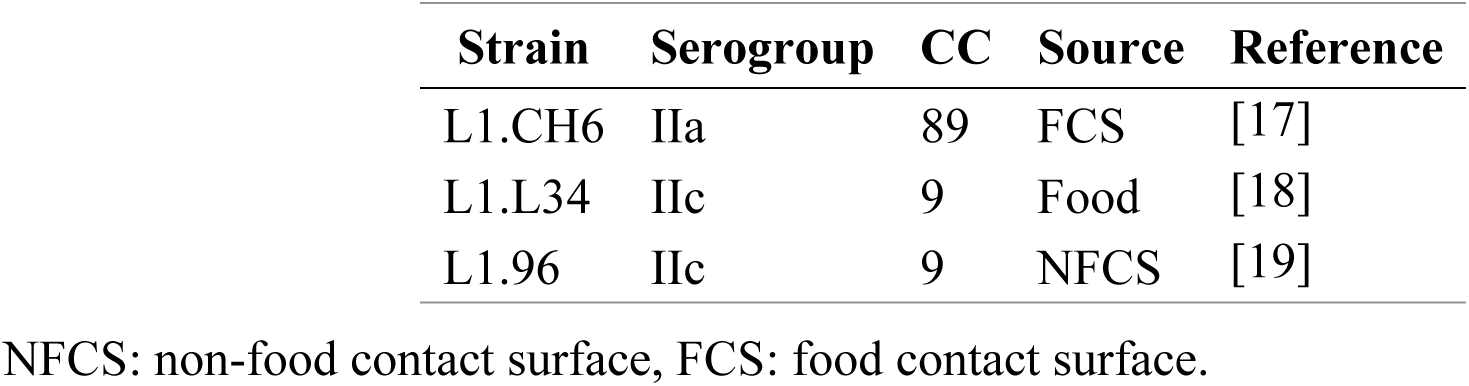
*L. monocytogenes* strains utilised in this study. Serogroups and clonal complex (CC) are shown for strain characterisation.

| Strain | Serogroup | CC | Source | Reference |
| --- | --- | --- | --- | --- |
| L1.CH6 | IIa | 89 | FCS | [17] |
| L1.L34 | IIc | 9 | Food | [18] |
| L1.96 | IIc | 9 | NFCS | [19] |

NFCS: non-food contact surface, FCS: food contact surface.

### 2.2. Experimental design

Inocula was prepared by adjusting reactivated cultures in fresh TSB to an OD_700_ of 0.1 ± 0.01, corresponding to approximately 10^8^ CFU/mL of TSB, as determined by previous calibrations. Concentrations were checked by plating in Tryptone Soy Agar (TSA; Scharlab, Barcelona, Spain), the plates were incubated at 37 °C for 24 h.

The biofilm and bulk cultures were prepared on 50x50x1 mm SS sterilised coupons placed in plastic petri dishes, containing 25 mL of 10^4^ CFU/mL TSB. Planktonic cell cultures were prepared in 100 mL culture flasks containing 25 mL of the same inoculum. Biofilms were cultured in moisture chambers under static conditions, and planktonic cells were cultured in shaking conditions (125 rpm); both systems were cultured at 25 °C for 72 or 120 h.

Next, planktonic cells were harvested by collecting a 5 mL aliquot from each flask. Bulk cells were recovered by collecting a 5 mL aliquot from the liquid phase of a biofilm. Next, the biofilms were washed twice with 25 mL of PBS to remove loosely attached cells. To ensure sufficient biofilm cell yield, the adhered biomass of 10 samples was pooled. Specifically, samples were sonicated 2 min at 40 Hz and aseptically scraped with a sterile plastic Digralsky sptula for 1 min in 10 mL of Phosphate-buffered saline PBS and RNA stabilisation solution (1:1; NZYTech, Lisboa, Portugal). Thus, the attached cells from 10 biofilms were concentrated to approximately 10 mL, then centrifuged to yield a 5 mL aliquot. The aliquots were spun for 5 min at 4 °C and 8000 *g*, the supernatant discarded, and resuspended in 5 mL of PBS and RNA stabilisation solution (1:1, NZYTech, Lisboa, Portugal). The aliquots were stored at 4 °C until RNA extraction. This experiment was performed in triplicate.

### 2.3. RNA sample preparation

For RNA isolation, the 5 mL aliquot was spun for 10 min at 4 °C and 8000 *g*, and the resulting pellet was resuspended in 100 µL of 15 mg/mL lysozyme solution, and incubated at 37 °C for 30 min. Next, RNA extraction was carried out using a NYZ Total RNA isolation kit (NZYTech, Lisboa, Portugal). Quantification of the resulting eluted RNA was performed in a Qubit™ 4 Fluorometer (Invitrogen, CA, USA) using the Qubit RNA HS Assay Kit (Invitrogen, CA, USA). RNA purity was checked with the 260/230 and 260/280 ratios, calculated in NanoDrop^TM^ 2000 (ThermoFisher, MA, USA). RNA quality was deemed acceptable if the ratios were 260/230 > 1.7 and 260/280 > 1.8. Finally, the RNAs were stored at –80 °C until use.

### 2.4. cDNA synthesis and reverse transcription quantitative PCR (RT-qPCR)

Complementary DNA (cDNA) synthesis was performed according to the NZY First–Strand cDNA Synthesis Kit (NZYTech, Lisboa, Portugal) using the manufacturer’s protocol. For each sample, a non–RT control was included. The reverse transcription (RT) reaction was performed in a MyCycler Thermal Cycler (BioRad, CA, USA) using the following conditions: a 10 min priming step at 25 °C, a 15 min reverse transcription step at 55 °C, and a 2 min RT inactivation step at 85 °C. As with RNA, cDNA concentrations were measured using the Qubit with the dsDNA HS Assay Kit (Invitrogen, CA, USA), and purity ratios were calculated using the NanoDrop. All resulting cDNAs were stored at – 20 °C until use.

The targeted genes for RT-qPCR are displayed in **Table 2**. The cDNA template was used in qPCR along with the iTaq Universal SYBR Green Supermix (BioRad, CA, USA). The mixture reaction contained 2 µL of cDNA, 1 µL of each forward and reverse primer (10 µM), and 10 µL iTaq Supermix, for a total volume of 20 µL. For each biological replicates, three technical replicates were done. The program consisted of an initial denaturing step of 30 s at 95 °C, followed by 40 cycles of 10 s denaturation at 95 °C and 1 min annealing and elongation at 60 °C. A final step of melting curve analysis was added by incubating the mix 15 s at 95 °C, followed by 1 min at 60 °C and 1 s at 95 °C. The program was run on a CFX Opus Dx Real–Time PCR (BioRad, CA, USA). The threshold cycle (Ct) was calculated using the program’s algorithm.

**Table 2.**
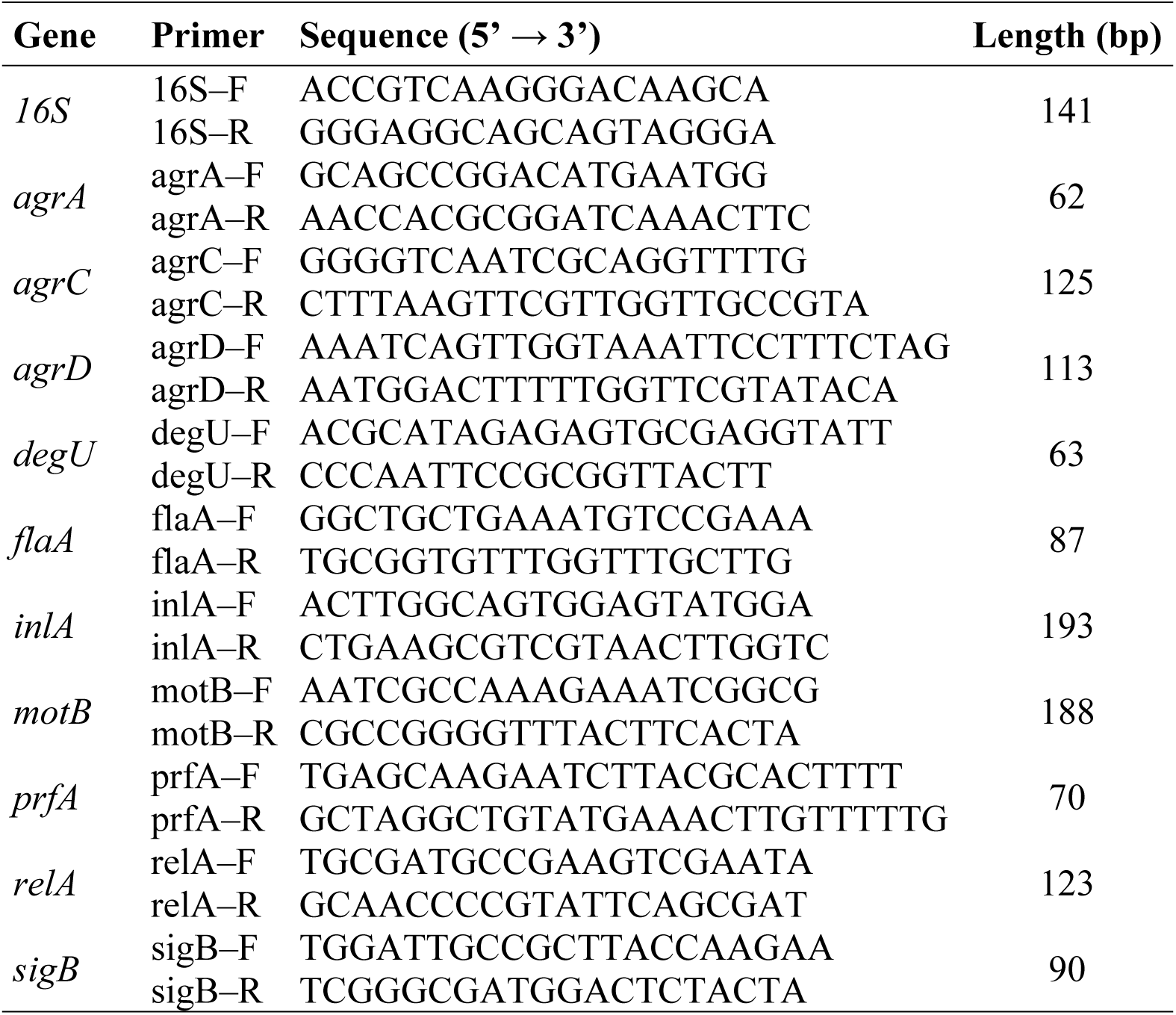
Primer sequences used for targeted genes on RT–qPCR assays. The primer sequences are taken from Liu et al. [20]. Length indicates the amplicon size.

Differential gene expression was assessed using Fold Change (FC), as described by Schmittgen and Livak, (2008). Briefly, the Ct corresponding to the 16S rRNA was used as a reference (internal control) to calculate the threshold cycle variation (ΔCt = Ct_gen_ – mean Ct_16S_), where Ct_16S_ was the average of the technical replicates (n=3) of the reference and Ct_gene_ is the Ct value of the corresponding target gene. Next, the ΔΔCt calculates the differential gene expression of biofilm and bulk cells with the planktonic cells as reference (ΔΔCt = ΔCt_biofilm or bulk_ – ΔCt_planktonic_), where the ΔCt_planktonic_ was calculated as the average of the technical replicates (n=3). Then, the FC was determined according to the Schmittgen and Livak, (2008) formula. When FC < 1, the proposed correction (- 1/FC) was applied to facilitate interpretation of downregulation. Finally, the mean values and Standard Error (SE) were calculated from the biological replicates. All calculations were performed in R as indicated in **Section 2.8**.

### 2.5. Strain growth kinetics under osmotic stress

To assess the strain growth kinetics at various osmotic levels, the inoculum was standardised and diluted to a final concentration of 10^4^ CFU/mL in TSB with different NaCl concentrations (0.5, 2, 4, and 8 %, w/v). Then, 200 µL of each inoculum was placed in a 96-well plate (Corning, MA, USA), with three biological replicates per strain and medium and three technical replicates per biological replicate. The plate was incubated at 25 °C for 50 hours. OD600 readings were taken every 20 minutes, preceded by a 5-second shaking step, using a Synergy H1 multi-mode microplate reader (BioTek, Vermont, USA).

### 2.6. Food model experiments

In these experiments, TSA plates containing different NaCl concentrations (0.5, 2, 4, and 8 %, w/v) were used as a food-simulating model. Briefly, 100 µL of the corresponding planktonic culture was dispensed in a single drop into the TSA plates in duplicate. The biofilms were washed twice with 25 mL of PBS, and a 30-s contact was done with the agar surface. Subsequently, the TSA plates were vacuum-packed in plastic bags using an ALFA Force^TM^ vacuum sealer (ALFA, Spain) and stored at 5 °C until sampling. Following incubation, cells were harvested with a loop, resuspended in 100 µL PBS containing 1:1 RNA stabilisation solution, and stored at 5 °C until RNA isolation.

RNA extraction was performed as described in **Section 2.3**, with slight modifications. Briefly, 45 µL of lysozyme at 50 mg/mL was added to the stored cell suspension to achieve a final lysozyme concentration of 15 mg/mL. The following steps of RNA extraction were done according to the protocol of the NZY Total RNA Isolation kit (NZYTech, Lisboa, Portugal). Each assay was performed with two independent RNA samples (biological replicates).

### 2.7. Bacterial survival after exposure to a simulated gastric environment

For the simulated gastric environment (SGE) experiments, gastric medium was prepared according to Beumer et al. [22] (**Table 3**). The pH was adjusted with HCl, and non–sterilised compounds (lysozyme and pepsin) were filtered through a 0.22 µm filter.

**Table 3.**
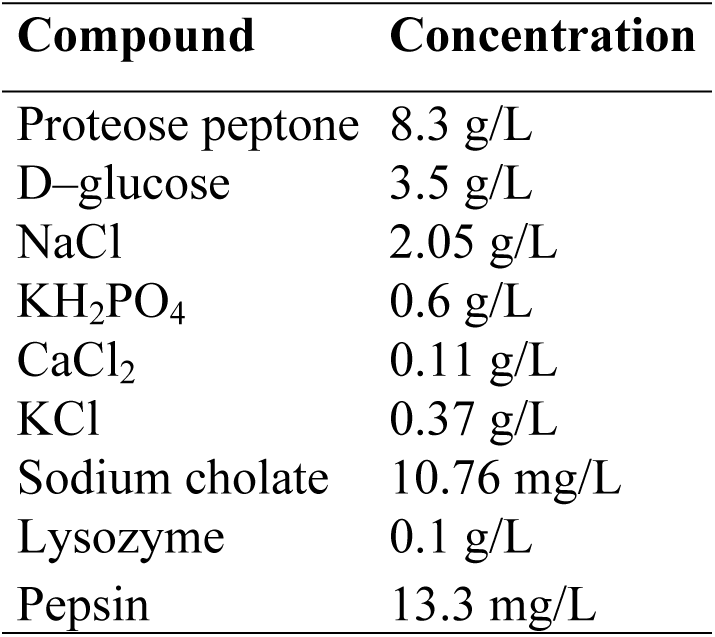
Simulated gastric environment composition.

Biofilm and planktonic cultures were prepared according to the procedures outlined in **Section 2.2**. After 72 h of culture at 25 °C, biofilm–attached and planktonic cells were transferred to the food system as detailed in **Section 2.6**. After 3 and 15 days of storage in the food model, cells were harvested in a microcentrifuge tube with a loop and resuspended in 200 µL of peptone water. The solution was adjusted to an OD_700_ = 0.1 ± 0.01 (108 CFU/mL) and serially diluted in the corresponding SGE to a final concentration of 10^5^ CFU/mL. Subsequently, 1 mL of the solution was added to a 24–well plate and incubated at 37 °C for 15, 30, and 60 min. At the selected time points, 100 µL aliquots were collected, plated on TSA, and cultured for 24 h at 37 °C for cell counting. For each pH range and cell state (biofilm–derived and planktonic–derived), the experiment was performed in triplicate.

### 2.8. Data analysis

Data analysis was conducted using R software version 4.3.0 [23] and RStudio version 4.5.1. [24]. All graphs were generated using *ggplot2* [25]. The survival-to-pH data were tested for normality and homocedasticity using the Shapiro-Wilk test and Levene’s test, respectively. Given the data structure and non-normality, a PERMANOVA was used to examine differences. We used the *adonis2* function from the vegan package (https://github.com/vegandevs/vegan) to examine the influence of cell type (biofilm-derived or planktonic-derived); osmotic stress (0.5 % NaCl and 4 % NaCl); TS, storage time (3 and 15 days); pH (1, 1.5, 3 and 7 pH); and the exposure to pH (15, 30, and 60 min) over the cell survival (log CFU/mL). The influence of the factors – cell type, osmotic stress, storage time and exposure to pH – was also analysed in each subset of pH. The pairwise comparison for the PERMANOVA was done with the *pairwiseAdonis::pairwise.adonis2* (https://github.com/pmartinezarbizu/pairwiseAdonis).

## 3. Results and Discussion

### 3.1. Differential expression of persistence and virulence genes: temporal dynamics and physiological state comparison

To investigate persistence and virulence-associated traits, the relative expression of a selection genes (**Table 4**) was assessed in biofilm and bulk cells, using planktonic cells as a reference. Gene expression was evaluated at 72 and 120 h, providing insight into temporal shifts and strain–specific patterns during biofilm maturation. Across all strains, a consistent time–dependent pattern emerged: biofilm cells generally showed upregulation at 72 h, followed by downregulation at 120 h, compared with their planktonic counterparts (**Figure 1**). This shift indicates that biofilm development initially correlates with increased activity of the selected genes, which decreases as the biofilm matures. In contrast, bulk cells exhibited highly heterogeneous profiles, showing simultaneous up- and downregulations of different genes at 72 h. This trend shifted toward a general downregulation at 120 h. However, a notable strain–dependent variability was found in the regulatory response, highlighting differences in the timing and magnitude of gene activation.

**Figure 1.**
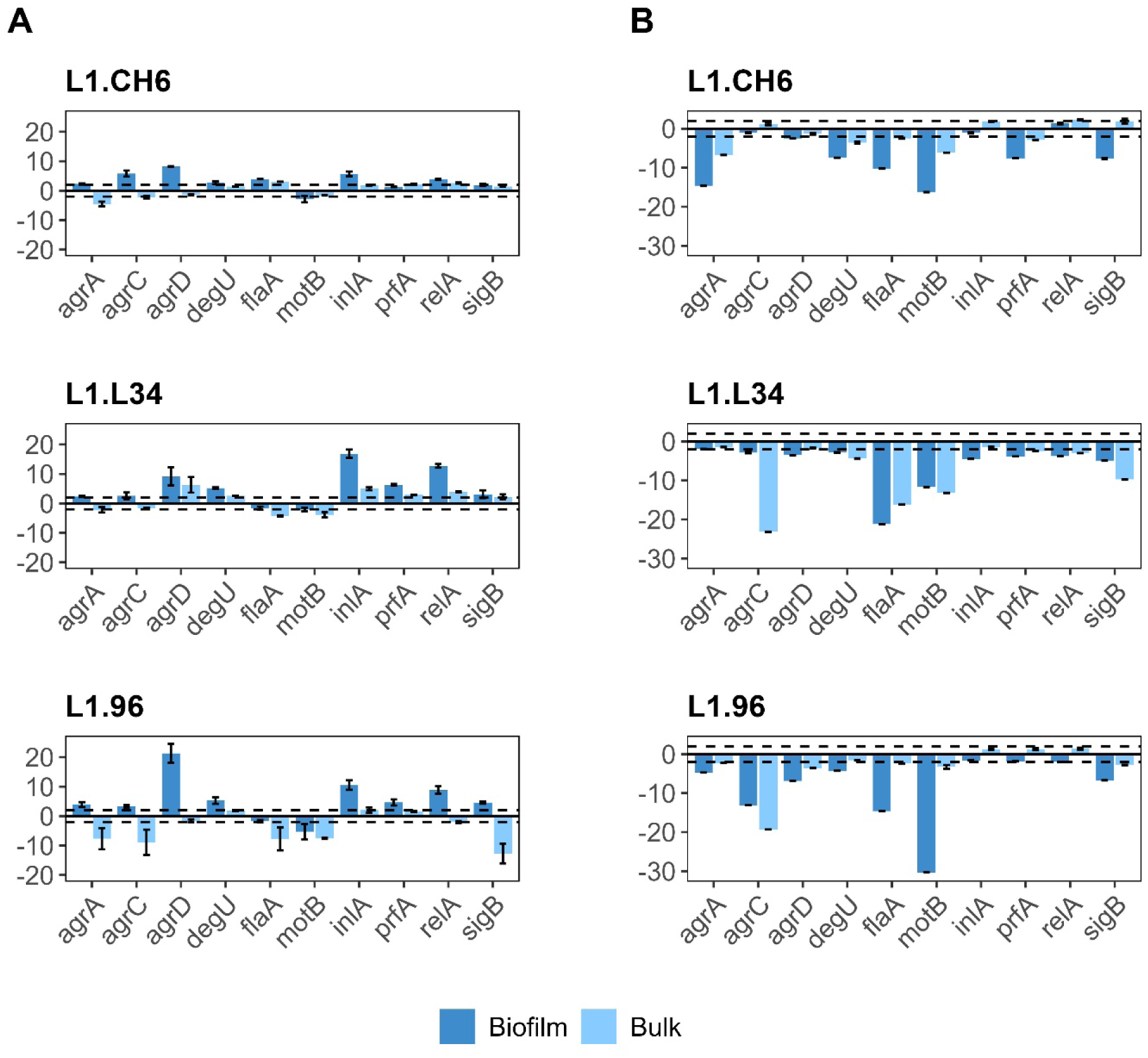
Differential gene expression in biofilm and bulk cells at different stages of biofilm development. **A**, gene expression at 72 h (early mature biofilm). **B**, gene expression at 120 h (late mature biofilm). Fold change (FC) values represent the expression in biofilm and bulk cells relative to that in planktonic cells. Data is presented as mean with standard error (n = 3). Dashed lines indicate the ± 2–fold change threshold for biological significance.

**Table 4.** Targeted genes used in this study.

| Gene | Locus tag<sup>1</sup> | Product | Process |
| --- | --- | --- | --- |
| *agrA* | lmo0051 | Response regulator | Quorum sensing |
| *agrC* | lmo0050 | Histidine kinase | Quorum sensing |
| *agrD* | lmo0049 | Auto-inducing peptide | Quorum sensing |
| *degU* | lmo2515 | Transcriptional regulator | Motility and stress response |
| *flaA* | lmo0690 | Flagellin | Motility |
| *motB* | lmo0686 | Flagellar motor rotation peptide motB | Motility |
| *inlA* | lmo0433 | Internalin A | Virulence |
| *prfA* | lmo0200 | Lysteriolysin positive regulatory protein | Virulence |
| *relA* | lmo1523 | Phosphokinase | Stress response |
| *sigB* | lmo0895 | RNA polymerase sigma factor B | Stress response |

<sup>1</sup> Locus tags from Elfmann et al. [26].

Specifically, at 72 h, biofilm cells from all strains showed increased expression of the *agr* quorum–sensing system (*agr*A, *agr*C, *agr*D), which regulates adhesion to abiotic surfaces, biofilm formation and virulence [27,28]. Among these genes, *agr*D was notably upregulated in L1.CH6 (8.22 ± 0.08 FC), L1.L34 (9.17 ± 3.10 FC), and particularly in L1.96 (21.29 ± 3.18 FC). This gene encodes the mature auto–inducing peptide that regulates the cell surface, and is thus required for cell adhesion [29]. Likewise, *deg*U, a pleiotropic transcriptional regulator involved in motility, biofilm formation and stress response [30,31], was upregulated in all biofilms. The upregulation was most pronounced in L1.L34 (5.21 ± 0.23 FC) and L1.96 (5.29 ± 1.11 FC), while L1.CH6 showed a modest increase (2.68 ± 0.49 FC), just above the 2–fold threshold.

Strain–specific differences were also observed in motility-associated genes at 72 h. Only L1.CH6 biofilm cells showed an increased expression of *fl*A (3.93 ± 0.02 FC), whereas levels remained below the threshold value in L1.L34 and L1.96. This was complemented by the downregulation of *mot*B observed in L1.CH6. While *fl*A encodes for flagellin [32] – a key component of the flagellum – *mot*B encodes a stator protein necessary for flagellar rotation and biofilm formation [33]. The simultaneous upregulation of *fl*A and downregulation of *mot*B suggests temporal regulation aimed at retaining the flagellum for stable adhesion while limiting its rotation. Since flagella are essential for stable attachment, the upregulation in L1.CH6 suggests that this strain maintains flagellar synthesis during the early maturation phase, potentially to facilitate further surface colonisation or to stabilise the biofilm structure. In contrast, the lack of differences in L1.L34 and L1.96 could indicate an earlier transition towards a more sessile state in these biofilms.

Virulence-related genes also showed a notable expression pattern in biofilm cells. A major point of strain-specific variability centred on *prf*A, a key virulence transcriptional regulator also implicated in biofilm formation and maturation [34,35]. At 72 h, L1.L34 (6.30 ± 0.26 FC) and L1.96 (4.62 ± 1.04 FC) displayed a clear upregulation compared to the planktonic counterparts, whereas L1.CH6 remained below the threshold. A similar yet distinct pattern was observed for the internalin gene *inl*A, which is regulated by *prf*A. While this gene was strongly upregulated in L1.L34 (16.85 ± 1.41 FC) and L1.96 (10.52 ± 1.61 FC) in coordination with *prf*A, L1.CH6 also exhibited a moderate upregulation (5.71 ± 0.79 FC) despite a lack of *prf*A activation. This suggests that *inl*A expression in L1.CH6 may be driven by alternative regulators, such as *sig*B, during early maturation [36,37]. This finding contrasts with previous studies reporting *inl*A downregulation in 24 h [38] and 48 h [39] biofilms, but aligns with a later study [40] showing significant upregulation in 144 h-old biofilms. Such discrepancies highlight both strain-specific variability and the dynamic nature of virulence gene expression during biofilm maturation.

Regarding the stress response, the regulators *rel*A and *sig*B showed similar strain-dependent expression patterns. *rel*A, which mediates the stringent response [41] – a survival mechanism triggered by stress, particularly starvation – was notably upregulated in L1.L34 (12.71 ± 0.61 FC) and L1.96 (8.85 ± 1.25 FC), whereas L1.CH6 showed a modest upregulation (3.87 ± 0.12 FC). The general stress regulator *sig*B mirrored this trend, with upregulations in L1.96 (4.56 ± 0.36 FC) and L1.L34 (3.12 ± 1.32 FC), while remaining below the threshold in L1.CH6. These results align with previous studies [10] reporting *sig*B upregulation in static biofilms compared to planktonic cells. The coordination between virulence and stress response genes suggests that the biofilms formed by L1.L34 and L1.96 are highly active reservoirs of virulent and stress–resistant cells.

By 120 h, a general strain-dependent downregulation of most target genes was observed in biofilm cells across all strains. This was particularly pronounced for motility-related genes. Thus, *fla*A and *mot*B were significantly downregulated across all strains. Specifically, *fla*A was reported to be 10-fold, 21-fold and 14-fold downregulated in L1.CH6, L1.L34 and L1.96, respectively. Likewise, *mot*B exhibited decreases of 16-fold, 11-fold and 30-fold in L1.CH6, L1.L34 and L1.96, respectively. This downregulation reflects a transition to a mature, quiescent biofilm state, where the energetic cost of maintaining the flagellar machinery is minimised. The transition from the slight upregulation (in L1.CH6) or no differences (in L1.L34 and L1.96) at 72 h to this profound downregulation at 120 h highlights a programmed shutdown of motility to prioritise metabolic efficiency and biofilm maintenance.

Similarly, *sig*B levels were downregulated across all strains, ranging from 4-fold in L1.L34 to 7-fold in L1.CH6. These findings are consistent with the reduced metabolic activity and global transcriptional downregulation typically observed in highly mature biofilms [42,43], indicating a widespread shift toward metabolic quiescence. However, some discrepancies exist in the literature. For instance, Toliopoulos and Giaouris, (2023) observed upregulation of *inl*A, *prf*A, and *sig*B in 144 h-old biofilms compared to their planktonic counterparts. These differences highlight the high phenotypic plasticity of *L. monocytogenes*, underscoring that regulatory responses are not only strain-specific but also sensitive to maturation stage and environmental conditions. The maturity–driven downregulation underscores that the biofilm state poses an evolving hazard, challenging the accuracy of traditional QMRA models and the efficacy of current monitoring strategies. Moreover, traditional *L. monocytogenes* detection methods based on metabolic activity may fail to capture quiescent cells within mature biofilms, leading to a false sense of safety. This is particularly concerning in established reservoirs, which could represent a heightened risk due to their potential for increased tolerance to sanitation. Thu, these results highlight the need for sanitation strategies that specifically target quiescent cells within mature biofilms.

In the bulk state, the differential gene expression reflects the transitional state of bulk cells. The intermediate nature of bulk cells is likely due to their heterogeneous composition of newly attaching, detached, and free-swimming cells. While bulk-state responses followed the trends observed in biofilms for a subset of target genes, the magnitude of the fold changes was generally lower. Specifically, at 72 h, no genes were notably upregulated in bulk cells. In contrast, L1.96 exhibited a notable downregulation of *agr*A, *agr*C, *sig*B, *fl*A, and *mot*B. By 120 h, no genes remained upregulated in the bulk state across any strain, and the downregulations intensified, particularly for *agr*C, *fl*A, *mot*B, and *sig*B in L1.L34, as well as *agr*C in L1.96. Notably, the downregulation of these genes (excluding *fl*A) was even more pronounced in bulk cells than in biofilms. This suggests that they may enter into deeper metabolic dormancy or transcriptional silencing than those within biofilms. This comparison emphasises that the cellular environment – whether a biofilm or the bulk state – dictates distinct, strain-specific regulatory outcomes for survival and persistence.

These findings support the hypothesis that the biofilm environment induces a more coherent and robust regulatory response than the transitional bulk state. Furthermore, these expression patterns characterise bulk cells as a unique physiological intermediate rather than a simple planktonic state. Evidence from other species supports this transitional nature. For instance, Rollet et al. [44] and Liu et al. [45] demonstrated that biofilm-detached cells of *Pseudomonas aeruginosa* and *Streptococcus mutants*, respectively, exhibit distinct phenotypes and transcriptomic profiles compared to both biofilm and planktonic cells. Similarly, Guilhem et al. [46] reported unique transcriptional signatures in biofilm-detached cells of *Klebsiella pneumoniae*. Consequently, bulk cells represent a critical link in the colonisation cycle, as they may either revert to a planktonic form or settle as new biofilm cells.

Collectively, the gene expression patterns indicate that 72 h-old biofilms maintain an active metabolism characterised by a strain-dependent upregulation of virulence genes, such as *prf*A and *inl*A, and reduced motility. This upregulation suggests a survival advantage and enhanced fitness for biofilm cells – likely mediated by *sig*B –, which not only promotes persistence but also increases the risk of higher *L. monocytogenes* loads in contaminated food products. Building on this evidence, it is critical to determine whether this enhanced fitness translates into differential stress tolerance in cross-contamination scenarios. This prompted a focus on *sig*B, the general stress regulator that mediates the bacterial response to environmental cues. In the saprophytic form, *L. monocytogenes* relies heavily on this factor for environmental adaptation. For instance, mutants lacking *sig*B exhibit impaired growth in soil compared to the wild–type strains [47]. Moreover, *sig*B is linked to virulence [48,49], by interacting with the *prf*A P2 promoter, where it directly regulates transcription [6]. Therefore, the subsequent section compares *sig*B expression in biofilm cells versus planktonic cells under a simulated cross-contamination scenario followed by cold storage, using this multi-strain approach to account for the observed phenotypic variability.

### 3.2. Differential expression of *sig*B in biofilm-derived and planktonic-derived cells under osmotic stress during food storage

The ability of *L. monocytogenes* to proliferate under a wide range of osmotic pressure conditions (up to 10-12 % NaCl) represents a major concern for food safety [50]. The halotolerance, combined with its psychrotrophic nature, allows it to colonise and grow in diverse RTE food products. Since food contamination by this pathogen typically originates from biofilm reservoirs rather than planktonic cultures, understanding how this prior cellular history influences subsequent adaptation to the food matrix is crucial. Therefore, to examine whether the physiological state – planktonic or biofilm – affects the stress response of *L. monocytogenes* during food storage, *sig*B expression was analysed in cells from both states following transfer to a TSA-based food model supplemented with 0.5, 2, 4, or 8 % NaCl (w/v). Food model units were vacuum-packed and stored under refrigeration, conditions well-documented to permit *L. monocytogenes* growth [51,52], and *sig*B expression levels were determined after various days of storage. This longitudinal analysis enabled assessment of both the immediate transcriptional impact of cellular origin and the long-term adaptive response to sustained osmotic stress under refrigeration.

Agar-based food models have been validated as surrogates for solid food matrices [53,54]. By eliminating the inherent biological variability of real food matrices, this approach ensures high reproducibility, allowing for a precise assessment of the osmotic stress response. Indeed, this reliability has been supported by reports of results comparable to those obtained in food products such as smoked salmon [54]. Furthermore, this model provides the flexibility to mimic a wide range of osmotic stress levels commonly encountered in RTE food products. This is particularly relevant considering that, while some products like cold-smoked salmon maintain water phase salt (WPS) levels between 3-4 % (w/w) [55], other processing techniques can result in considerably higher concentrations, as seen in salt–cured fish or salted anchovies [56]. Since 8 % NaCl has been reported to impair *L. monocytogenes* [57], the growth dynamics of the three strains were first monitored in liquid cultures to ensure that the selected osmotic stress levels did not compromise cellular viability. Kinetic monitoring confirmed that cells remained viable and metabolically active across all salt concentrations studied. These conditions were therefore suitable for assessing the *sig*B response (**Figure 2**). Furthermore, these growth profiles reveal strain-specific physiological differences, providing additional context for the transcriptional patterns discussed below.

**Figure 2.**
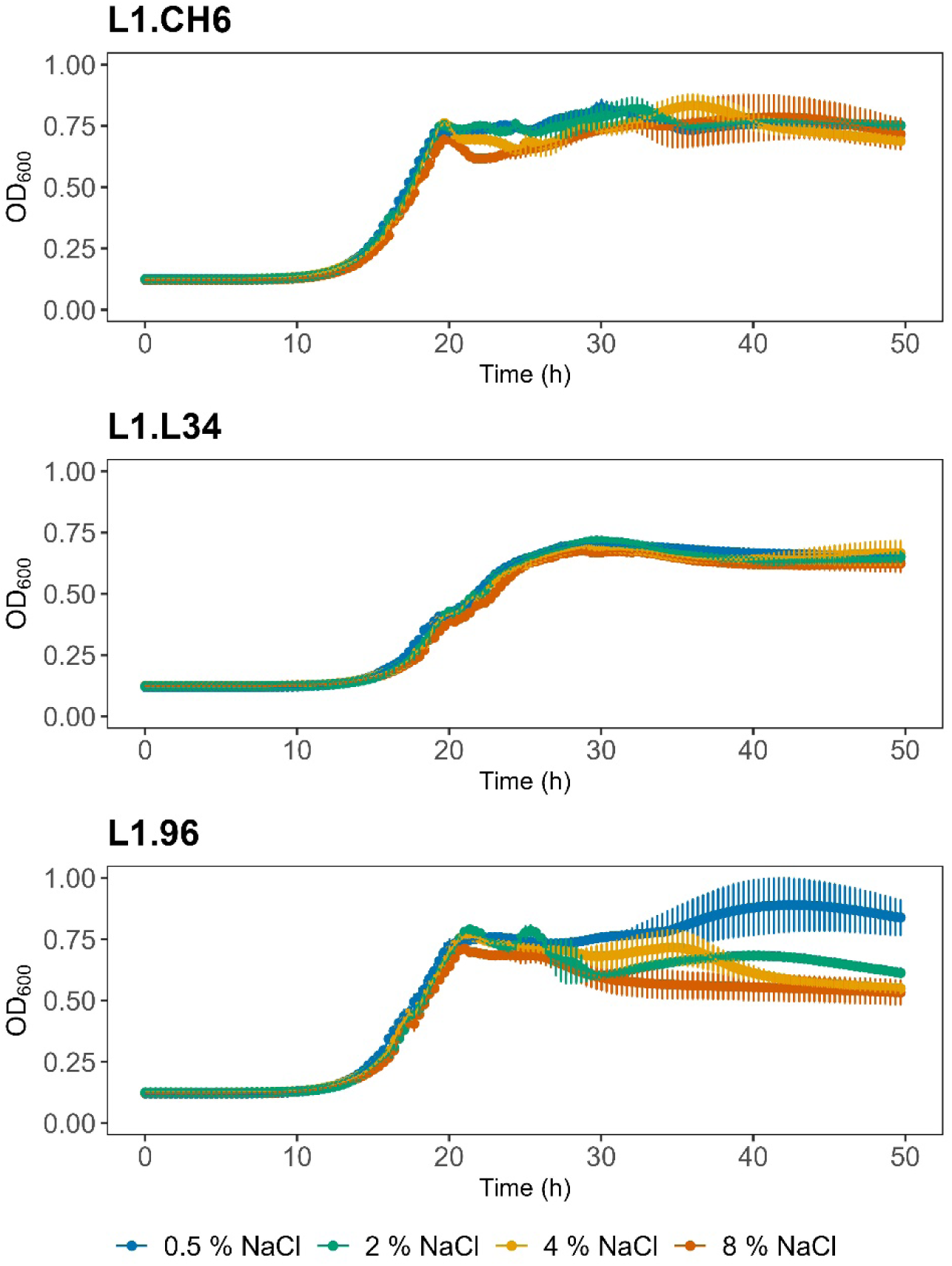
Strain’s growth kinetics under different osmotic stresses. The growth dynamics in liquid cultures were measured at 20–minute intervals for 50 hours. Data represent mean values and standard error (n = 3).

#### 3.2.1. Comparative analysis across three *L. monocytogenes* strains: impact of cellular origin

The differential expression levels of *sig*B during storage are presented in **Figure 3**. On day 5 of storage, strain L1.96 showed a marked upregulation of biofilm-derived cells relative to planktonic cells across all salinity levels. This response followed a dose-dependent trend, peaking at 4 % NaCl (13.53 ± 1.56 FC), with notable increases also observed at 2 % NaCl (9.91 ± 4.34 FC) and 0.5 % NaCl (7.05 ± 0.07 FC). However, under severe stress (8 % NaCl), this differential expression diminished notably (4.54 ± 0.67 FC). This suggests that *sig*B expression in biofilm- and planktonic-derived cells converges in both populations under high salinity, attenuating the influence of prior physiological state. This trend aligns with findings by Utratna et al., [58] in liquid cultures, where *sig*B expression increased proportionally with NaCl concentration (0.5-5.3 %).

**Figure 3.**
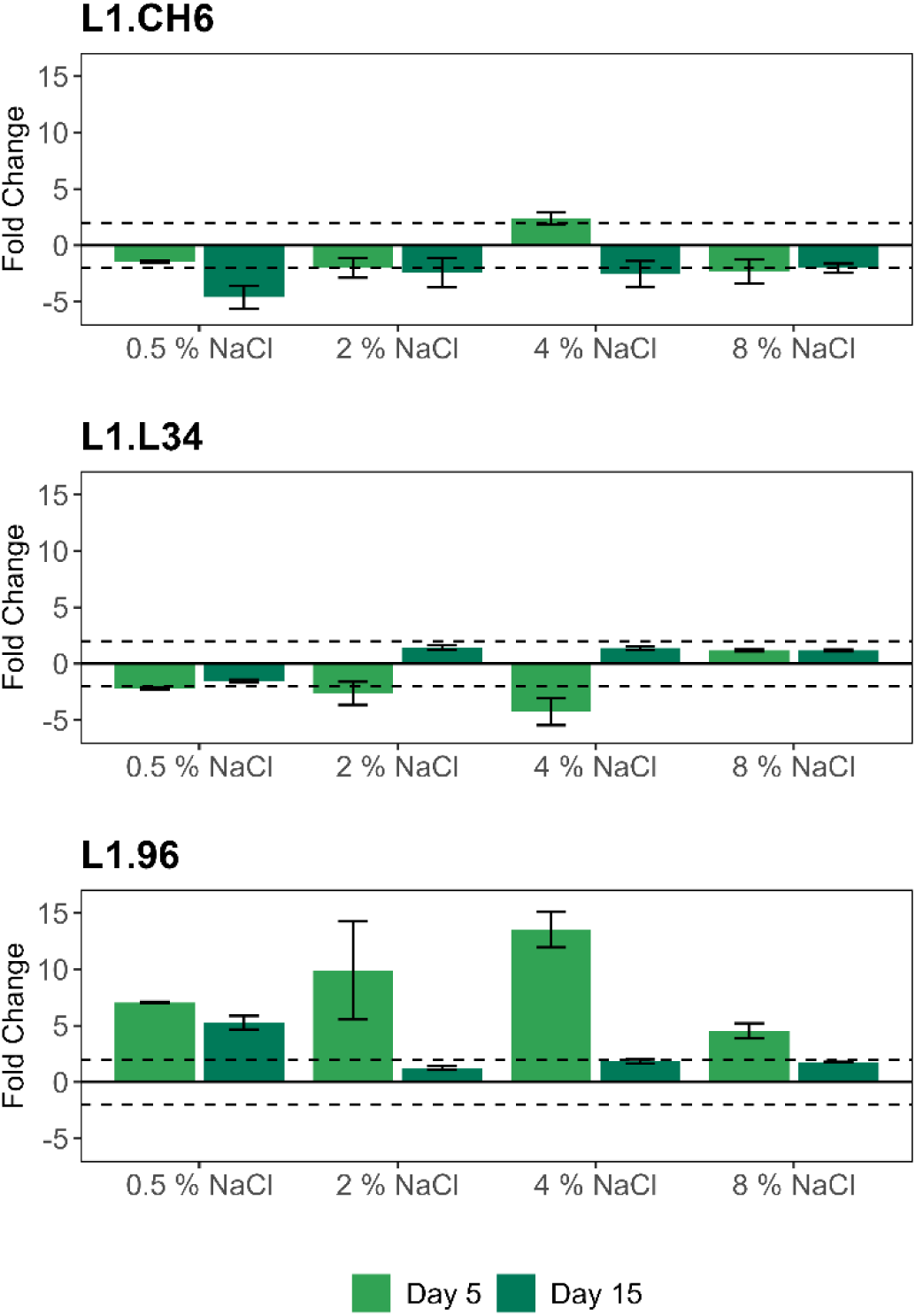
Differential *sig*B expression in biofilm–derived cells under osmotic stress during food-model storage. Fold change (FC) values represent *sig*B expression in biofilm-derived cells relative to planktonic-derived cells. Data is presented as mean with standard error (n = 2). Dashed lines indicate the ± 2-fold change threshold for biological significance.

Conversely, biofilm-derived cells of L1.L34 and L1.CH6 showed minimal differential expression at day 5 of storage, with FC values fluctuating around the 2-fold threshold across several concentrations. Specifically, in L1.CH6, levels remained near the threshold under all conditions: 0.5 % NaCl (-1.45 ± 0.08 FC), 2 % NaCl (-2.00 ± 0.88 FC), 4 % NaCl (2.37 ± 0.53 FC), and 8 % NaCl (-2.31 ± 1.06 FC). Similarly, L1.L34 exhibited responses near the threshold at 0.5 % (-2.23 ± 0.19 FC) and 2 % (-2.62 ± 1.02 FC), whereas a consistent downregulation was observed at 4 % NaCl (-4.24 ± 0.10 FC). These results suggest minor fluctuations rather than a coordinated differential stress response at this early stage.

These findings reveal a clear strain-dependent differential response in biofilm-derived cells at 5 days of osmotic stress exposure. Specifically, the marked upregulation in L1.96 at day 5 may suggest that biofilm-derived cells of this strain activate a specific stress response upon exposure to the simulated food matrix and storage conditions. This is consistent with the prior cellular history, where 72 h-old biofilm cells already showed *sig*B upregulation (**Figure 1**). This enhanced stress-response profile appeared to be maintained in subsequent generations, rendering them pre-adapted to osmotic stress upon transfer. This phenomenon, known as bacterial memory [59,60], refers to the ability of bacteria to retain information about past physiological states, thereby modulating future gene expression and phenotypic responses [61].

The existence of bacterial memory in biofilm cells has been explored by Miyaue et al. [62], who demonstrated that past environmental exposures can shape subsequent stress tolerance. Specifically, they observed that colony-biofilm cells of various gram-negative (*Escherichia coli*, *Acinetobacter*, and *Salmonella*) and gram-positive pathogens (*Bacillus* and *Staphylococcus*) retained higher antibiotic tolerance than their planktonic counterparts, even after biofilm dispersal. The authors hypothesise that the memory effect may result from a specific molecular signalling pathway established during biofilm maturation. They also suggest that cell dormancy within biofilms may last longer after biofilm withdrawal, enabling cells to maintain the persister phenotype. This memory effect can arise from epigenetic [63] and metabolic [64] mechanisms, such as DNA methylation or sustained protein expression, thereby enhancing cellular fitness in new environmental conditions.

By day 15 of storage, biofilm-derived cells of all three strains generally exhibited minimal to no differential *sig*B expression. In the case of L1.L34 and L1.CH6, this lack of response reflected a high degree of stability in *sig*B expression, allowing them to handle osmotic stress with minimal shifts throughout storage. Regardless of their prior lifestyle (biofilm or planktonic), these strains maintain consistent expression levels in the food model without requiring further compensatory induction. Nonetheless, minor exceptions were observed and, thus, L1.L34 and L1.CH6 displayed slight downregulation under 4 % NaCl at day 5 and 0.5 % NaCl at day 15, respectively. In contrast, for L1.96, the increased expression profile inherited from biofilm cells appeared to fade over successive generations. According to Wolf et al. [61], who exposed *Bacillus subtilis* to sporulation medium containing antibiotics, the short-term memory (transient) is prevalent over long-term memory. Therefore, bacterial memory is predominantly subject to a timescale limit [59]. Then, most physiological adaptations from past environments dissipate over time through successive generations, eventually leading the population toward greater homogeneity.

This time-dependent decay explains why, in the present study, the *sig*B expression levels of planktonic- and biofilm-derived cells in L1.96 became nearly indistinguishable by day 15 under most conditions. As both populations adapt, they transition from a state dominated by their prior history towards a converged physiological state in which the influence of the biofilm origin is eventually superseded by prevailing environmental pressures. Nevertheless, this convergence did not occur at 0.5 % NaCl, where biofilm-derived cells still maintained upregulation (5.27 ± 0.60 FC). Although the limited number of biological replicates (n = 2) requires cautious interpretation, this suggests that the biofilm memory can persist longer under low-stress conditions, keeping the two populations distinct.

Beyond validating metabolic activity under osmotic stress as established earlier, the growth kinetics of all three strains in liquid culture (**Figure 2**) provide further context for these differential expression patterns and serve as an indicator of intrinsic physiological sensitivity. Although L1.96 reached the highest population levels under low-stress conditions (0.5 % NaCl), and showed a faster growth rate than L1.L34 across all salt levels, it proved to be the most sensitive to salinity, exhibiting a marked decline in cell density during the stationary phase at 2-8 % NaCl. This suggests a biological trade-off where L1.96 prioritises rapid growth in optimal environments at the expense of robustness under stress.

Under the restrictive conditions of the study (biofilms and vacuum storage at 5 °C), the intrinsic sensitivity of L1.96 is likely further exacerbated. Biofilms impose severe restrictions – nutrient gradients, oxygen limitation, and the accumulation of metabolic products – which probably trigger greater metabolic stress in L1.96 than in L1.L34 and L1.CH6 (more robust strains). Consequently, the high FC values of biofilm-derived cells in L1.96 at day 5 might reflect both the persistence of a stress-response profile and a pronounced compensatory induction aimed at restoring cellular equilibrium in the food model. In contrast, for L1.L34 and L1.CH6, the distinction between biofilm and planktonic origin was less critical. Their intrinsic resilience was reflected in the absence of a marked differential transcriptional response between the two cellular origins. This suggests that their adaptive capacity is an intrinsic trait of such strains rather than a condition-dependent advantage, indicating that they did not require a primed state from a biofilm to effectively activate their stress-response mechanisms. Consequently, the safety of RTE foods is challenged by a dual phenomenon: the history-dependent readiness of biofilm-derived cells from sensitive strains, in which cellular origin determines the magnitude of the initial response, and the history-independent resilience of robust strains, which maintains equivalent *sigB* profiles regardless of their prior environment.

#### 3.2.2. Time-dependent differential *sig*B expression in strain L1.96

While the initial comparative study provided a snapshot of the osmotic stress response at key stages of storage, the unique transcriptional behaviour of L1.96 prompted a more detailed kinetic analysis. To achieve this, a second experimental stage was conducted, focusing exclusively on this strain. By integrating new time points (1, 3, and 10 days) with the initial data (5 and 15 days), a comprehensive kinetic profile of the *sig*B response was established. This high-resolution approach allowed for the characterisation of rapid shifts that might otherwise have been missed in the broader comparative analysis, providing a deeper understanding of how cellular origin dictates the speed and magnitude of stress acclimation.

As shown in **Figure 4**, results exhibited a clear temporal trend: differential expression peaked at different time points within the first 5 days, depending on salt concentration, and then declined at later stages (days 10 and 15). At 0.5 % NaCl, upregulation was highest during the first three days (14.02 ± 2.95 FC and 13.95 ± 2.73 FC), and decreased consistently thereafter, falling to 5-fold by day 15. However, this pattern shifted under higher salt concentrations. At 4 % and 8 % NaCl, the initial response on day 1 (9.03 ± 1.96 FC and 2.66 ± 0.63 FC, respectively) was lower than on day 3 (15.20 ± 3.38 FC and 5.06 ± 0.22, respectively), after which expression declined steadily. A similar profile was observed at 2% NaCl, although the peak was delayed until day 5. Despite high variability between replicates at this point, their values remained higher than at any other time point, confirming a differential *sig*B expression peak. By days 10 and 15, expression at 2 % NaCl fell below all previous levels. Indeed, considerable variability was observed among replicates across various conditions. The variability is likely attributable to stochastic gene expression and cell population heterogeneity, as described by Guldimann et al. [65]. And would be avoided by increasing the number of replicates and time points.

**Figure 4.**
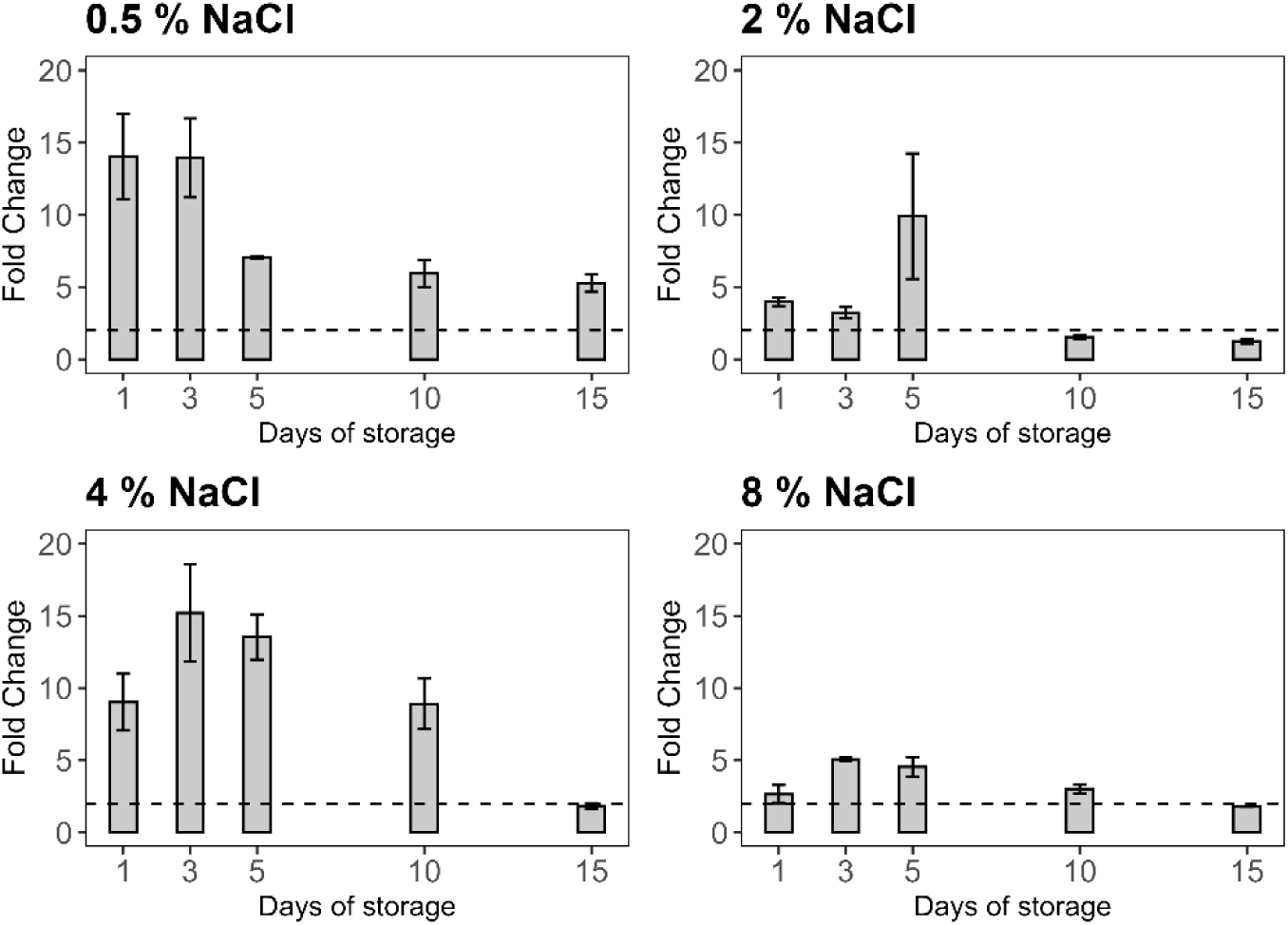
Time–dependent *sig*B expression in biofilm–derived cells under osmotic stress during food–model storage. Fold change (FC) values represent *sigB* expression in biofilm–derived cells relative to planktonic–derived cells. Data is presented as mean with standard error (n = 2 or n = 3). Dashed lines indicate the ± 2–fold change threshold for biological significance.

Substantial differences were also observed across salinity levels. Differential expression was generally higher at 4 % NaCl than at 2 % or 8 %, except on day 15, when levels became comparable and fell below the threshold. Notably, the response at 0.5 % NaCl was consistently higher than at 2 % and 8 %, except on day 5, when levels at 0.5 % and 2 % were similar. Conversely, the response at 4 % NaCl frequently surpassed that of 0.5 %, particularly on days 5 and 10. Collectively, these findings confirm a dynamic, time–dependent differential *sig*B response in L1.96 biofilm-derived cells, characterised by a strong upregulation during the early stages of exposure, particularly under moderate salt conditions. This early activation is likely triggered by the need to promptly activate the metabolic machinery for growth in the new environment. A recent meta-analysis has underscored the role of *sig*B in regulating carbon metabolism, ion transport, and protein metabolism [6], supporting the idea that it is essential for the initial physiological transition. However, this differential response diminished over time as both cell types converged toward a common physiological state. By the later stages of storage, the *sig*B profile is thus no longer dictated by cellular origin, but by the requirements of the current environment.

These kinetic profiles have direct implications for the safety of RTE products. The fact that *sig*B upregulation in biofilm-derived cells peaked within the first 5 days and remained higher than in their planktonic counterparts for up to 10 days – particularly at 0.5 % and 4 % NaCl – suggests a significant window of risk during food distribution. This advantage was even more persistent under low-stress conditions (0.5 % NaCl), where differential expression was maintained up to day 15. Although this head start is less pronounced under extreme salinity (8 % NaCl), the overall trend indicates that, in a real-world scenario such as cold-smoked salmon (typically 3-6 % WPS), *L. monocytogenes* reaches its maximum adaptive state once the product is already moving through the distribution chain, and consumption is most likely. This timing is particularly concerning, as it effectively extends the window in which the pathogen may be better equipped to survive subsequent stresses, such as the low pH of the human stomach.

Overall, L1.96 biofilm-derived cells exhibited a *sig*B response that likely enhances cellular fitness by promoting metabolic adaptation and survival. These results demonstrate that the interplay between previous transcriptional states and current environmental pressure determines the adaptive trajectory of *L. monocytogenes*. While cellular origin provides a significant advantage in expression during the first days of storage, this influence is gradually replaced by a specialised response tailored to the severity of the osmotic challenge. However, while increased *sig*B expression suggests greater resilience, transcriptional changes do not always translate directly into a phenotype of increased virulence or survival. Therefore, it is necessary to validate whether this transcriptional readiness actually confers a survival advantage under stressful conditions, such as those encountered upon human consumption. To test this hypothesis, the stress tolerance of biofilm-derived cells was compared with that of planktonic-derived cells by exposing both populations to a simulated gastric environment to determine whether the *sig*B upregulation observed here confers enhanced survival under acute acid conditions. This approach provides direct phenotypic validation of the potential risk to consumers.

### 3.3. Acid tolerance of salt preconditioned biofilm-derived and planktonic-derived cells in a simulated gastric environment

After enduring cumulative stresses in RTE food products during storage, *L. monocytogenes* must withstand the extreme challenges of the human gastrointestinal tract. The first and most critical barrier in this environment is the extreme acidity of the stomach, where pH typically ranges from 1 to 2.5 [66]. A previous study demonstrated that *L. monocytogenes* can survive for up to 2 h in simulated gastric fluids [67], a critical timeframe for reaching the intestine and initiating infection. The adaptive response to acidic conditions within the host is well-documented as being *sig*B-dependent [4]. In fact, mutants lacking *sig*B have shown much lower acid survival rates than wild-type strains [68].

Consequently, phenotypic assays were conducted to determine whether the higher *sig*B expression in L1.96 biofilm-derived cells would be translated into a functional survival advantage during a subsequent gastric challenge. This assessment focused on strain L1.96, as it was the only strain that exhibited a marked *sig*B induction during food storage, providing a unique model to test whether this divergence translates into enhanced physiological fitness. Specifically, biofilm- and planktonic-derived populations were recovered from the food model under non-stress (0.5 % NaCl) or osmotic stress (4 % NaCl) conditions at specific time points (3 and 15 days). Then, cells were exposed to a simulated gastric environment (SGE) across a range of pH values (1-7) to assess cell survival. This multifactorial design was used to analyse how the interplay between cellular origin, osmotic conditions, and storage duration shapes the resilience of *L. monocytogenes* against lethal gastric acidity. Finally, PERMANOVA analyses were conducted to identify the main effects and interactions of these variables, examining both the entire dataset and each pH level separately.

As shown in **Figure 5**, survival varied greatly across pH levels. At SGE pH 7, cell counts remained stable under all conditions (p > 0.05). Similarly, at SGE pH 3, no significant differences (p > 0.05) were observed between biofilm- and planktonic-derived populations, and osmotic preconditioning had no effect. At this pH, the only significant reduction was observed in planktonic-derived cells preconditioned at 0.5 % NaCl after 3 days of storage, where cell counts decreased significantly (p < 0.05) after 60 min of exposure. However, survival dynamics changed significantly under extreme acidity, with a clear advantage observed for biofilm-derived cells at pH 1.5 and especially at pH 1, compared to their planktonic counterparts across several experimental conditions. Consistent with these observations, the initial PERMANOVA on the full dataset identified pH as the primary driver of mortality, accounting for 87 % of the total variability, with a large effect size (ω^2^ = 0.43). As pH accounted for most of the variance, separate PERMANOVA analyses were conducted for each pH level.

**Figure 5.**
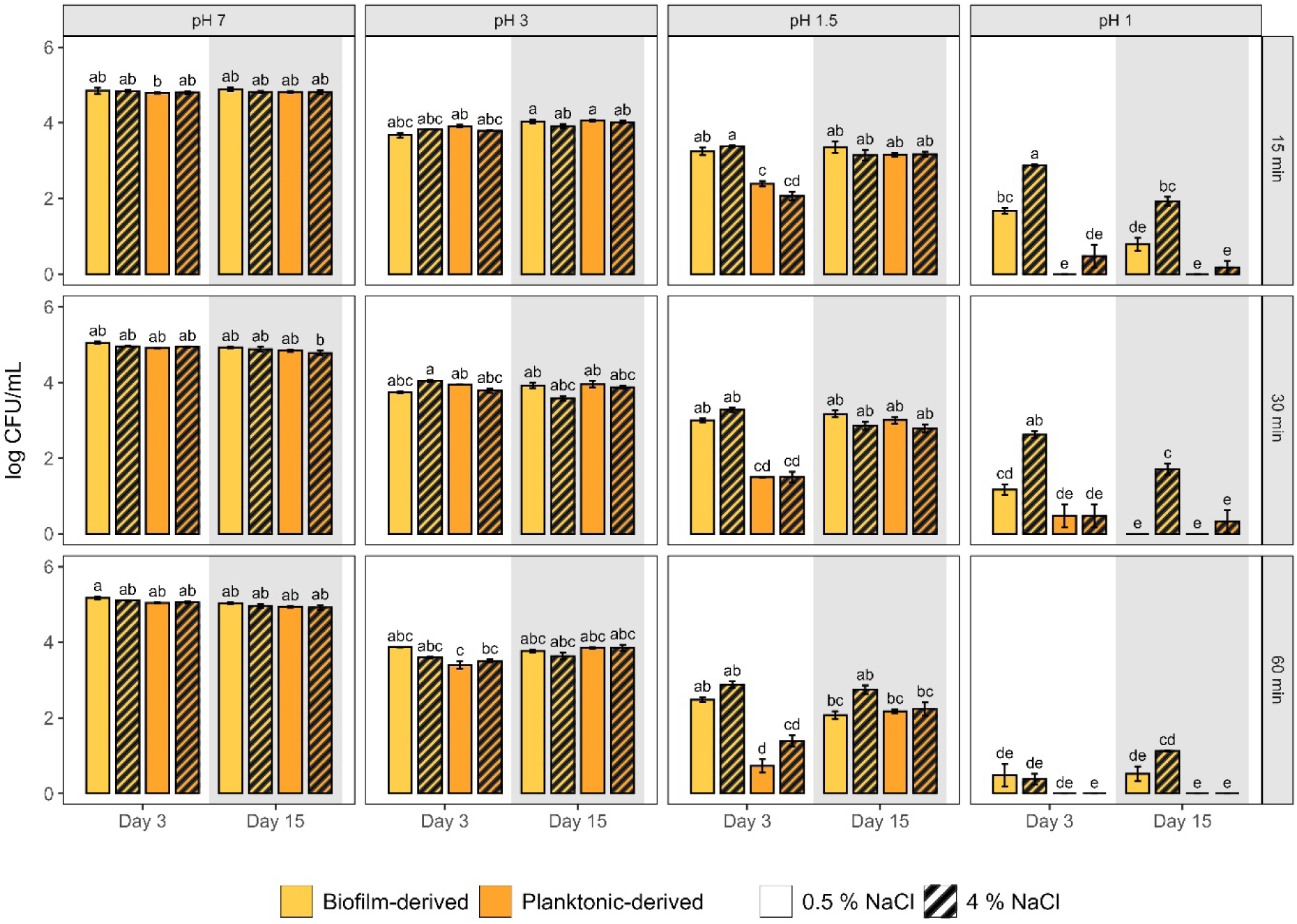
Survival of L1.96 biofilm- and planktonic-derived cells in a simulated gastric environment (SGE). Data is presented as mean with standard error (n = 3). Different letters indicate significant differences (p < 0.05) within each pH; no comparisons were made between different pH levels.

Within these individual models, cellular origin emerged as the most influential factor for survival under extreme acidity, explaining 44 % and 28 % of the variance at pH 1 and 1.5, respectively. These results showed large effect sizes (ω^2^ = 0.22 and ω^2^ = 0.14), confirming that the physiological legacy of biofilm origin is a critical determinant of acid tolerance rather than a minor effect, as further demonstrated by the high percentage of variance explained solely by cellular origin at pH 1 (44 %). Furthermore, the interaction of cell type, salt concentration, storage time, and pH exposure time also explained a substantial proportion of the total variance, amounting to 33 % (ω^2^ = 0.16) at pH 1.0 and 24 % (ω^2^ = 0.10) at pH 1.5. Finally, the individual effect of exposure time accounted for 9 % (ω^2^ = 0.05) and 25 % (ω^2^ = 0.12) of the variance at pH 1 and 1.5, respectively.

Specifically, at 3 days of storage, biofilm-derived cells exposed to SGE pH 1 for 15 min showed significantly higher counts than planktonic-derived cells (p < 0.05), regardless of the salt preconditioning level. However, at longer exposure times (30 and 60 min), viability declined sharply, and the biofilm-origin advantage persisted only for 30 min in populations preconditioned at 4 % NaCl. A similar survival advantage was observed at 15 days of storage, but only in cells preconditioned with 4 % NaCl, where this resilience remained significant across all three exposure times (p < 0.05). For instance, after 15 min of exposure, survival was significantly higher in biofilm–derived cells than in planktonic counterparts (1.93 ± 0.11 log CFU/mL *versus* 0.18 ± 0.18 log CFU/mL; p < 0.05). In contrast, no significant differences (p > 0.05) were found between biofilm- and planktonic-derived cells preconditioned at 0.5 % NaCl. Notably, regardless of the storage time, a binary survival pattern emerged in populations exposed for 60 min at SGE pH 1. While planktonic-derived cells were consistently reduced below the detection limit, biofilm-derived cells maintained a resilient residual population across nearly all tested conditions. Specifically, from an initial load of approximately 5 log CFU/mL, a surviving fraction of around 1 log CFU/mL was recorded at the end of the challenge. This underscores that, under sustained extreme acidity (60 min), the survival advantage of biofilm-origin cells becomes an exclusive trait.

At SGE pH 1.5, survival was higher than at pH 1 across several of the experimental conditions. Similar to the findings at pH 1, a clear survival advantage for biofilm–derived cells was observed after 3 days of storage under both salt conditions. However, in this case, the advantage persisted across all exposure times. This gap closed after 15 days of storage, partly due to increased resilience of planktonic cells, which exhibited higher survival than at day 3. Notably, salt concentration significantly affected the survival of biofilm-derived cells at pH 1 but not at pH 1.5. Salt-induced cross-protection was evident during the early stages of exposure (15-30 min), with survival significantly higher in cells preconditioned with 4.0 % NaCl than with 0.5 % NaCl (p < 0.05). For instance, after 15 min, counts reached 2.88 ± 0.03 log CFU/mL and 1.67 ± 0.08 log CFU/mL, respectively. In contrast, this protection was not observed in their planktonic counterparts.

Clinical observations of the human gastric environment in the fasted state have reported that pH frequently drops to 1-2.5 [66]. Under these conditions, *L. monocytogenes* survival becomes critically dependent on the glutamate decarboxylase and the arginine deiminase systems [69]. Given that the *sigB* regulon directly governs their expression [6], osmotic stress likely pre-activates the adaptive machinery. This allows biofilm-derived cells to reach the gastric barrier with their defences already functional, a response that appears less efficient in planktonic counterparts. Despite its significance, to the authors’ knowledge, the literature has only briefly addressed differences in survival between biofilm- and planktonic-associated *L. monocytogenes* phenotypes under gastric conditions.

On one hand, a recent study by Serrano-Heredia et al. [70], also found that biofilm-derived cells on cooked ham exhibited higher survival during in vitro digestion (at pH 2 and 3) than their planktonic counterparts immediately after transfer. However, they found that this gap narrowed significantly after a week of storage, suggesting that planktonic cells may develop a biofilm-like phenotype during storage. The present findings at pH 1.5 reinforce this hypothesis, as the difference between biofilm- and planktonic-derived cells eventually disappeared after 15 days of storage. This suggests that identical conditions in the food matrix trigger phenotypic convergence, in which planktonic cells appear to catch up to the stress-resistance levels of their biofilm-derived counterparts. Nevertheless, it is crucial to note that this equalisation is not absolute. While storage conditions may level the survival under milder acidic stress, the extreme challenge of SGE pH 1 continues to reveal a superior, residual fitness in biofilm-derived cells throughout the entire experimental period. This implies that while both cells may develop a similar phenotype during storage, the biofilm legacy confers a maximum threshold of resistance that planktonic cells – even after adapting to the food matrix – cannot fully match when facing the most severe environmental frontiers.

On the other hand, Bai et al. [39] reported no significant differences in survival based on cellular origin in two strains tested after exposure to simulated gastric fluid at pH 2. This discrepancy is noteworthy, as both Serrano-Heredia et al. [70] and the results reported in this study found differences at that pH level or slightly lower. This discrepancy may be explained by three factors: first, the biofilm legacy is a strain-specific phenotype rather than a universal trait. Notably, Serrano-Heredia et al. [70], also utilised L1.96 among the strains they tested. Secondly, the intensity of the acidic challenge and the narrow threshold for survival are decisive. The current research has identified that as acidity increases, the survival landscape for *L. monocytogenes* can shift dramatically. Even a small shift in pH (from 2 to 1.5) may reveal resilience advantages that are less evident at higher pH levels, potentially favouring the viability of a larger population and thus increasing the probability of successful infection. Third, regarding the method used to dislodge cells, this study and Serrano-Heredia et al. [70], transferred biofilm cells by contact to food, while Bai et al. [39] used sonication to detach them. Although sonication is effective for recovering cells, it might alter their physiological state developed during biofilm development, thereby masking the survival advantage observed under contact-transfer conditions.

On balance, three main effects can be highlighted, starting with a significant survival advantage for biofilm-derived cells at SGE pH 1 and 1.5, regardless of the NaCl conditions. Additionally, there is a salt-induced enhancement in the survival of biofilm-derived cells at SGE pH 1; a phenomenon of heterologous cross-protection previously described in *L. monocytogenes* [71]. For instance, pre-exposure to 6 % NaCl has been shown to upregulate *sigB*-dependent genes, providing cross-protection against oxidative stress [72]. Lastly, a storage-dependent effect at SGE pH 1.5, where the advantage of biofilm-derived cells vanished by day 15. These findings demonstrate a fitness advantage acquired during biofilm development, likely mediated by *sig*B upregulation, which enables a rapid acid-stress response. This is further amplified by salt pre-conditioning, which acts as a potent activator of the *sig*B regulon, as previously shown.

This response aligns with the concept of bacterial memory, suggesting that this advantage persists through the initial stages of food storage. As observed by Miyaue et al. [62], biofilm cells can retain their antibiotic resistance for several weeks after dispersal from the biofilm. However, the present findings suggest that bacterial memory is a transient legacy that is progressively lost, eventually levelling the fitness between cell origins. This is evidenced by a dual physiological process where planktonic cells gradually adapt to the food matrix – as seen in their improved survival at SGE pH 1.5 –, while biofilm-derived cells undergo a gradual resetting of the stress-response machinery, supported by the previously reported decrease in *sigB* upregulation over time (**Figure 4**).

Nevertheless, the present findings at SGE pH 1 reveal that this convergence remains incomplete within the tested period. Even when both populations were subjected to identical sustained osmotic stress within the food model, their physiological backgrounds remained distinct, suggesting that the conditions acquired in biofilms imprint a resilience that adaptation to a nutrient–rich food matrix does not fully replicate. Consequently, while planktonic cells can narrow the fitness gap under acidic stress at SGE pH 1.5, the biofilm origin confers a superior threshold of protection that remains the ultimate determinant of survival under extreme challenge. Whether these differences would eventually vanish over extended storage beyond 15 days remains to be elucidated.

Moreover, the survival of biofilm-derived cells after a 60-minute challenge at SGE pH 1 highlights a significant extension of the survival window compared to planktonic cells. Although the surviving fraction was small (∼ 1 log CFU/mL), this residual population can be biologically determinant. Such a binary survival pattern, in which only biofilm-derived populations maintain a resilient fraction, suggests that biofilm origin may promote phenotypic heterogeneity, potentially leading to the emergence of persister-like cells. It remains to be elucidated, however, whether such cells could withstand a longer gastric challenge to further define the limits of this biofilm-mediated advantage.

These findings have relevant implications for food safety management, particularly regarding the timing and site of contamination within food processing environments. In many RTE food products - such as cured meats or cold-smoked fish - where a final lethal treatment is absent, contamination can occur at any stage of the production chain. If contamination originates from biofilms established on abiotic surfaces -such as conveyor belts or slicers-, *L. monocytogenes* enters the food supply chain already primed for extreme acid gastric resistance. Unlike cells of planktonic origin, biofilm-derived cells, which remain highly vulnerable to gastric conditions, biofilm-derived cells possess an intrinsic robustness that makes them immediately hazardous upon ingestion. Consequently, maintaining rigorous hygienic design and specialised biofilm control throughout the entire processing line is essential, as these cells may reach the consumer before their specialised physiological state can be attenuated. This underscores that the risk associated with contamination events is defined not only by the bacterial load but also by the physiological history of the cells at the point of entry into the food.

Further implications arise when consumer preferences are considered. Labels such as “Use-by date” and “best-before” dates strongly influence consumer choices, with willingness to pay and product acceptance dropping significantly as expiration approaches [73]. This preference highlights a neglected dimension of food safety that aligns dangerously with the biofilm legacy. The fact that biofilm-derived cells are inherently more resilient than their planktonic counterparts -coupled with their peak resilience during the initial stages of shelf life-may effectively lower the infectious dose threshold precisely when consumption is most likely. These phenomena render environmental history a critical determinant that can even turn a low-level contamination event into a successful infection. Consequently, the present evidence suggests that this physiological background must be integrated alongside strain characteristics and contaminating loads as a key input in quantitative microbial risk assessment models. Nonetheless, while this study demonstrates a survival advantage provided by the biofilm origin, further research is required to explore whether this advantage translates into increased *in vivo* virulence. Moreover, it remains to be elucidated how the duration of the bacterial memory varies across different RTE food categories, and whether the loss of fitness differs between growth-supporting and non-growth-supporting food products.

## 4. Conclusions

In conclusion, this study offers new insights into the differences in fitness and survival of *L. monocytogenes* biofilm-derived cells compared with their planktonic counterparts, raising concerns about the potential underestimation of risks when only planktonic cultures are used in food safety assessments. Therefore, incorporating the biofilm’s physiological background into risk assessment frameworks is essential for developing accurate control strategies against *L. monocytogenes* in FPE. Ignoring this biofilm legacy may lead to a systemic underestimation of the resilience of *L. monocytogenes*.

Overall, the evidence presented in this study provides valuable insights for food safety management and QMRA, while opening interesting paths for further study. For instance, further research is required to explore whether the biofilm-derived advantage translates into increased in vivo virulence, it also remains to be elucidated how the duration of the bacterial memory varies across different RTE food categories, and whether the loss of fitness differs between growth–supporting and non–growth–supporting food products. Translating these laboratory insights into the complexity of FPE requires a broader ecological perspective.

## Author contributions statement (CRediT)

R.A.N. was responsible for the investigation, formal analysis, and writing of the original draft. J.J.R.H. provided conceptualisation and supervision, contributed to the writing of the original draft and was involved in funding acquisition. P.R.L. contributed to the methodology and investigation. M.L.C. provided conceptualisation and supervision, contributed to the writing of the original draft and was involved in funding acquisition. All authors reviewed and edited the manuscript.

## Acknowledgements

The authors thank S. Rodríguez-Carrera for her valuable support with all the experimental work. R. Nogueira expresses her gratitude to Dr C. O’Byrne and Dr J. Wu for their helpful support and thoughtful advice. P. Rodríguez-López gratefully acknowledges Universidad San Jorge (USJ) for the financial support through the USJ Postdoctoral Fellowship Programme.

## Funding declaration and data statement

This study was funded by the Spanish Ministry of Science and Innovation (ASEQURA, PID2019-10 8420RB-C31, ASEQURA2, PID2023-152192OB-C21) and the Spanish Research Council (PIE 202170E074). The funders played no role in study design, data collection, analysis and interpretation of data, or writing of this manuscript.

Data is available upon request.

